# Temporal regulation of TRP channels during partial sciatic nerve ligation is modulated by PLC□ in male mice

**DOI:** 10.64898/2026.03.02.709140

**Authors:** Beatriz C. de Moraes, Alexandre M. do Nascimento, Elaine F. Toniolo, Ana M. Rodrigues, Carolina P. Goes, Camila S. Dale, Deborah Schechtman

## Abstract

Neuropathic pain is a debilitating condition afflicting millions worldwide, still lacking a proper and effective treatment. Understanding the underlying mechanisms that lead to neuropathic pain may lead to the discovery of new targets. Previously, we found that PLC□ is a key player in mechanical hypersensitivity triggered by either capsaicin or complete Freund’s adjuvant induced inflammation. Here, we investigated the role of PLCγ in neuropathic pain using a partial sciatic nerve ligation (PSNL) injury model in male mice and a peptide inhibitor of PLCγ activation (TAT-pQYP) injected at different time points after injury. Mechanical hypersensitivity was reversed by TAT-pQYP at 7, 14, and 28 days post-injury (dpi). Furthermore, both TAT-pQYP and the TRPA1 inhibitor HC-030031, but not the TRPV1 inhibitor capsazepine, were able to reverse mechanical hypersensitivity at 14 dpi. In contrast, all three treatments significantly improved the mechanical threshold at 28 dpi. After a challenge with TRPV1 agonist capsaicin at 14 dpi, TAT-pQYP treated animals exhibited increased sensitivity. We also found decreased TRPV1 mRNA levels in the DRG of PSNL animals at 14 dpi that returned to baseline at 28 dpi. TAT-pQYP treatment normalized TRPV1 expression to sham levels at 14 days. Conversely, TRPA1 mRNA expression increased at 14 days and returned to baseline at 28 days, while TAT-pQYP reduced TRPA1 expression at 14 days. At this same time point, injection of a pan-PLC (U73122), or Trk (GNF-5837) inhibitor reduced mechanical hypersensitivity and decreased *Trpa1* expression, whereas only PLC□ inhibition by TAT-pQYP or Trk inhibition restored *Trpv1* levels to Sham values. These findings indicate that PLCγ, and Trk signaling are important for expression of *Trpa1* and activity of *Trpv1*. Taken together, our results show that dynamic changes in *Trpv1* expression in the PSNL model account for differential sensitivity to TRPV1 inhibition and suggest that clinically relevant pharmacological inhibitors targeting ion channels may vary in efficacy depending on the temporal regulation of expression during neuropathic pain development. Finally, we show that TAT-pQYP is able to reverse mechanical hypersensitivity arising from neuropathic pain, adding evidence that PLCγ is a promising target to be explored for neuropathic pain management.

**Significance statement:** We show that the efficacy of inhibiting TRPV1 or TRPA1 on neuropathic pain in a partial sciatic nerve ligation model and that pain mitigation depends on the dynamic changes of channel expression post-injury, in the dorsal root ganglion, emphasizing that temporal regulation of drug targets should be taken into account when choosing therapeutic strategies for pain treatment. Notably, we identify PLCγ as a promising therapeutic target, as its inhibition with TAT-pQYP reverses mechanical hypersensitivity during different stages of injury progression by modulating expression and activity of TRP channels.

## INTRODUCTION

Neuropathic pain, resulting from nerve injury or dysfunction, is one of the most challenging forms of pain to manage, characterized by heightened sensitivity to mechanical stimuli and involving complex molecular signaling pathways within both central and peripheral nervous systems^1^. Despite significant advances, the precise mechanisms that underpin neuropathic pain remain poorly understood, underscoring the urgent need for novel, more effective analgesic therapies^2,3^.

Amongst the key players leading to neuropathic pain in the peripheral nervous system are the transient receptor potential (TRP) cation channels, which mediate nociceptive signaling in response to a variety of direct and indirect stimuli, of note vanilloid receptor 1 (TRPV1) and transient receptor potential ankyrin 1 (TRPA1), are the most prominent chemosensitive TRP channels in nociceptive neurons^4,5^, and are broadly explored as analgesic targets in both inflammatory^6^ and neuropathic pain^7^. TRPV1 is directly activated by noxious heat, low pH, vanilloids^8^, and indirectly by phospholipase C (PLC) cleavage of Phosphatidylinositol 4,5-bisphosphate (PIP2), that binds to the channel inhibiting its activity^9^. Furthermore, protein kinase C (PKC) activation downstream of PLC signaling leads to phosphorylation of TRPV1 potentiating channel activation^10^. In turn PLCγ activation is mediated by receptor tyrosine kinases (RTK), upon growth factor (GF) binding, and PLCβ by G-protein receptor signaling mediated by Gαq activation^11,12^. During inflammatory processes GF and activators of G-protein signaling are released by immune cells, leading to both PLC□ and β activation^9,13,14^. TRPA1 is directly activated by noxious cold, allicin, cinnamaldehydes and oxidized lipids^15–17^, and has emerged as an important analgesic target in the context of neuropathic pain since it is described as a sensor for oxidative stress^18^. As TRPV1^19^, inhibition of TRPA1 channel by lipids such as PIP2 has also been proposed^20^. Expression and activity of these two ion channels is frequently increased in neuropathic/chronic pain in somatosensory neurons^21–23^, and in some cases decreased^24,25^. These ion channels have also been described to physically interact with each other, suggesting a potential co-regulatory mechanism^26^. However, what regulates their expression in somatosensory neurons and whether there are changes in the kinetics of expression in different pain models is still under investigation, an important question that can help explain the variety of responses regarding the efficacy of treatments with modulators of these channels.

Previous work from our group showed that inhibiting PLCγ using a permeable phosphopeptide, TAT-pQYP, designed to inhibit the interaction between receptor tyrosine kinases (RTKs), such as Tropomyosin receptor kinases (Trks) and PLCγ, a key interaction for lipase activation, had an antinociceptive activity in two inflammatory models: in the acute and chronic phase of the adaptive immunity induced pain caused by Complete Freund’s Adjuvant (CFA) model of inflammatory pain^27^, and during capsaicin induced neurogenic inflammation^13^. In the present study, we explore the role of PLCγ in neuropathic pain induced by nerve injury after partial sciatic nerve ligation (PSNL) in male mice. A dynamic gene expression of TRP channels in the dorsal root ganglia (DRG) was observed during progression of the model. When compared to Sham operated animals, *Trpa1* was upregulated at 14 dpi, and normalized at 28 dpi, while *Trpv1* was downregulated at 14 dpi, and normalized at 28 dpi. Subcutaneous injection of TAT-pQYP in the hind paws of mice at 14 dpi, besides having an analgesic effect, decreased *Trpa1* levels and normalized *Trpv1*, restoring sensitivity to capsaicin. Taken together, we show that in the PSNL model in male mice, hypersensitivity at 14 dpi is mainly due to TRPA1, and that expression coding for TRPA1 and TRPV1 is dynamic, regulated by PLCγ and thus may explain differential sensitivity to channel inhibitors.

## MATERIALS AND METHODS

### Animals and Bioethics

Male C57BL/6NTac mice weighing 20–26 g, age-matched (4 to 9 weeks of age), were used for this study. Animals were maintained under controlled light cycles (12/12 h) and temperature (21±2 °C) with free access to food and tap water. Animal work was carried according to the ARRIVE 2.0 guidelines (Animal Research: Reporting of In Vivo Experiments)^28^. Research designs were approved by the Ethics Committee on the Use of Animals at University of São Paulo (CEUA, protocol No. 4121050423).

### Partial Sciatic Nerve Ligation

Peripheral neuropathic pain was induced in mice by PSNL as previously described^29^. Briefly, animals were deeply anesthetized with isoflurane (5% for induction and 2% for maintenance) in O_2_. The left sciatic nerve was exposed after the incision of skin and the dorsal 1/3 to 1/2 of the sciatic nerve was tightly ligated with an 8–0 silk suture just distal to the point at which the posterior biceps-semitendinosus nerve branches off. After, the muscle and skin layer were at once sutured and the animal was moved to a heating pad for anesthesia recovery.

### Pharmacological modulators and peptides

All drugs or substances were injected subcutaneously (s.c.) to the dorsal surface of the hind paw using 29 G needles in a 20 µL volume. TAT-pQYP (GRKKRRQRRRPQGSGQAPPVpYLDVLG) were diluted to 1.6 µmol/paw in distilled water. Peptide concentration and the 3 hour timepoint for behavior analysis was selected based on our previous studies^13,27^. Inhibitors were solubilized in DMSO, and diluted as follows: Capsazepine to 600 pmol/paw^30^; U-73122 to 100 pmol/paw^13,27^; GNF-5837 to 2 µmol/paw^13,27^; HC-030031 to 100µg/paw^31^, LY294002 to 10µg/paw^32^.

### Mechanical sensitivity assessment

To evaluate mechanical sensitivity, mice were placed individually in acrylic cages above an elevated wire mesh floor that allowed access to their paws. Mice were habituated to the experimental environment for 1h previous to any intervention. Von Frey Filaments (Touch-Test Sensory Evaluators, North Coast Medical, Morgan Hill, CA), ranging from 0.07 g to 6.00 g, were applied to the plantar region of the left hind paw of each animal for 5 seconds, and the response profile interpreted according to the up-down method^33^. Mechanical sensitivity measurements were performed before PSNL surgery (baseline - BSL) and at 7, 14 and 28 dpi, and 3h post-modulators injection

For mechanical sensitivity to innocuous touch, the 0.07 g monofilament was applied 6 consecutive times to the hind paw, in intervals of at least 30 seconds, as previously described^13,34^. Mice and treatments were randomized (AMdN) and the experimenter (BCM) was blinded to the treatments.

### Capsaicin test

At 14 dpi, mice were injected with modulators and challenged with capsaicin 2.00 µg/paw (s.c.) in 10% ethanol and 10% tween-20^30^. Immediately after injection of the irritant, the animals were placed in an acrylic apparatus positioned on top of absorbent matte paper and in front of a straight mirror for better visualization of the animal. The mice had their behavior recorded for 5 minutes and the videos were analyzed to quantify the latency of painful behavior (s), which consisted of licking or biting the paw ipsilateral to the injection^13^. Mice and treatments were randomized (BCM) and video analysis were conducted blindly (AMdN).

### RT-qPCR

Total RNA was isolated from L4, L5 and L6 DRGs using Trizol reagent (Invitrogen, Carlsbad, CA, USA). Briefly, total RNA (1 μg) was reverse-transcribed by oligo(dT) primer using a SuperScript IV cDNA Synthesis Kit (Thermo Scientific). This cDNA was used for qPCR (20 ng/reaction) using Power SYBR green Master mix (Thermo) with three technical replicates per biological replicate, using 200 nM of the primer sets, TRPV1: Forward - 5’ CACCCTGAGCTTCTCCCTG 3’, Reverse - 5’ TATCTCGAGTGCTTGCGTCC 3’; TRPA1: Forward - 5’ CAACTCACTCACTCCCACCT 3’, Reverse - 5’ TGGCTTCAAAGAGAGGGGAG 3’; ATF3: Forward - 5’ GCGCGGATCCATGATGCTTCAACATCCA 3’, Reverse - 5’ GCGCAAGCTTTTAGCTCTGCAATGTTCCTTC 3’; GAPDH: Forward - 5’ ACTTCAACAGCAACTCCCACT 3’, Reverse - 5’ TGGGTGGTCCAGGGTTTCTTA 3’. Reactions were performed on an Applied Biosystems 7500 Real-time PCR system, using default settings. GAPDH was used as housekeeping control and standard 2^−ΔΔCt^ analysis performed relative to the Sham control sample.

### Statistical analysis

Data was analyzed using GraphPad Prism (10.3.0). Raw data were assessed for normality by fitting to the normal distribution curve. Data in graphs are presented as mean□±□standard deviation (SD). p-values <□0.05 were considered statistically significant. A detailed description of the statistical test, post-hoc test, number of replicates per group and multiple comparison data are available in Supplementary data file S1.

## RESULTS

### TAT-pQYP reversed mechanical hypersensitivity induced by PSNL

TAT-pQYP antinociceptive activity in a neuropathic pain model induced by PSNL was assessed at 7, 14 and 28 dpi (Fig. 1A), and mechanical hypersensitivity was consistent with previous reports^35–37^, and reverted after intraplantar injection of a PLCγ inhibitor peptide, TAT-pQYP, at three hours post injection at all days tested (Fig. 1B). Injection of the peptide to Sham operated animals 14 days after surgery did not alter mechanical sensitivity (Fig. 1C). These findings corroborate our previous data on two different inflammatory models^13,27^, where the same concentration and time point after injection evaluated, was able to induce antinociception in challenged mice, without impacting naïve or sham animals. We also assessed the expression of the nerve injury marker, activating transcription factor 3 (ATF3)^22,38^, in the DRG (L4-L6) of operated mice at different time points (7, 14 and 28 days post-surgery), confirming that our model was able to elicit the transcriptional remodeling characteristic of sciatic nerve damage, and that TAT-pQYP treatment was unable to modulate this phenotype (Fig. 1D).

**Fig. 1:**
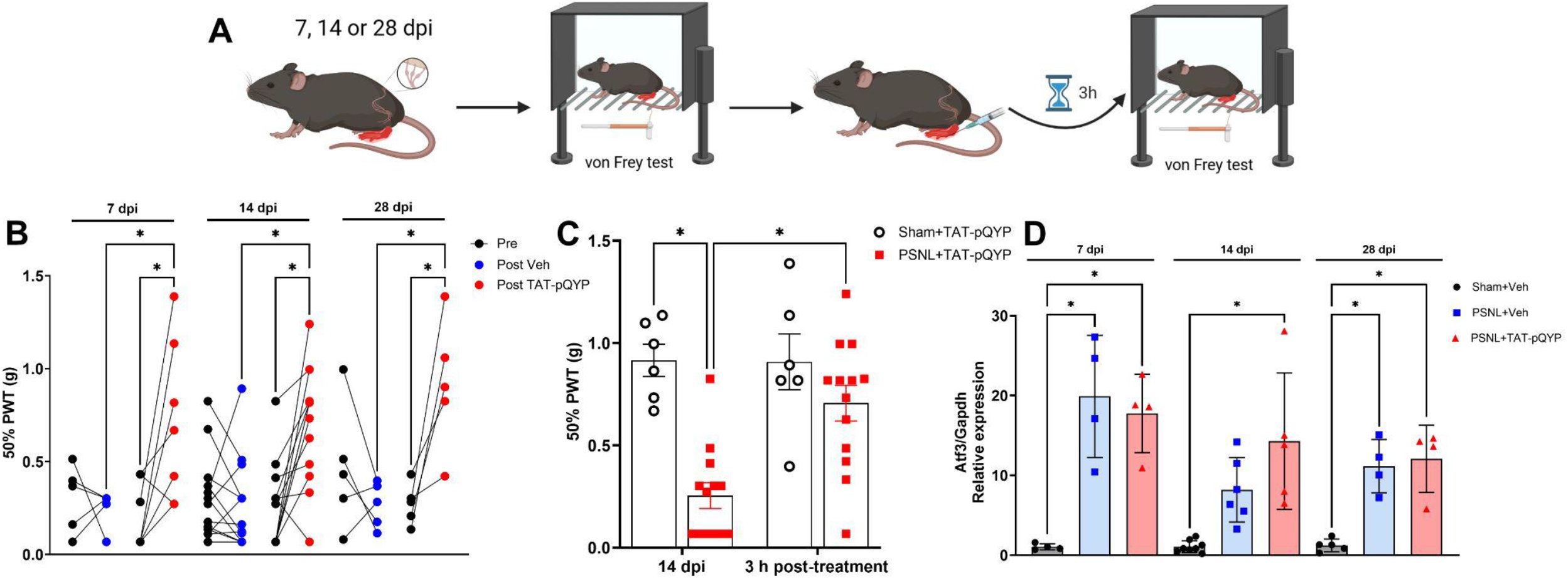
TAT-pQYP reverses mechanical hypersensitivity elicited by PSNL, but does not interfere with transcriptional remodeling induced by nerve lesion. (A) Experimental protocol adopted (figure created in BioRender.com). (B) TAT-pQYP reversed the PSNL mechanical threshold to baseline levels at 7, 14 and 28 dpi. (C) TAT-pQYP reverses mechanical hypersensitivity at 14 dpi. (D) ATF3 mRNA levels are upregulated after PSNL at 7, 14 and 28 dpi, and are not affected by TAT-pQYP treatment. Data are expressed as mean ± SD. One-Way ANOVA followed by Šídák’s (B, D) or Fisher’s (C) multiple comparison tests for significance. * stands for p < 0.05.

### Selective inhibition of TRPA1 and PLC□, but not TRPV1, reverses mechanical hypersensitivity after PSNL

To further assess the role of PLC□ and TRPA1/V1 channels in our model, and considering the limitation that PSNL often leads to the minimum threshold of our traditional up-down von Frey approach, we tested how the selective inhibition of these targets impacted the innocuous touch perception, which is exacerbated during pain^39^. TAT-pQYP was able to reverse the hypersensitivity to innocuous stimuli, similarly to TRPA1 inhibitor HC-030031 (Fig. 2A) at 14 dpi, while animals treated with a combination of the TRPV1 inhibitor (CPZ) and the TRPA1 inhibitor also showed reversal of the sensitivity threshold. However, TRPV1 inhibition alone had no effect, indicating that the increased nociceptive behavior observed at 14 dpi is dependent on TRPA1, but not on TRPV1 (Fig. 2A). However, at 28 dpi, either TAT-pQYP, CPZ and HC-030031 were able to reverse the hypersensitivity, indicating a possible temporal regulation of these channels (Fig. 2B).

**Fig. 2:**
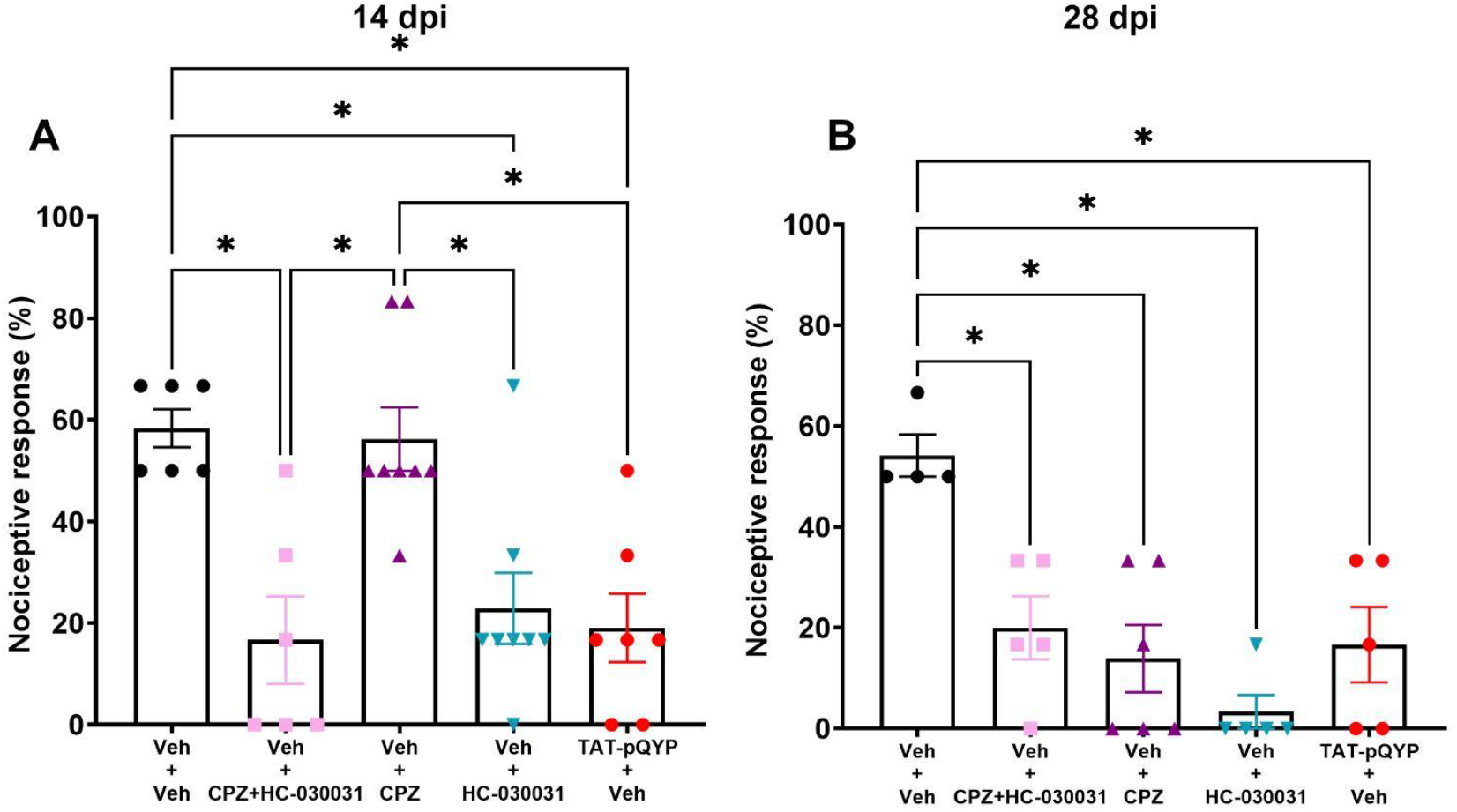
TRPV1 inhibition, opposed to TRPA1 and PLC□, fails to reverse hypersensitivity at 14 days post-injury. (A) TAT-pQYP and HC-030031 reverses mechanical hypersensitivity after innocuous stimuli, while capsazepine (CPZ) fails to modulate nociception at 14 dpi. (B) TAT-pQYP, HC-030031 and CPZ effectively reverses mechanical hypersensitivity to innocuous stimuli at 28 dpi. Data are expressed as mean ± SD. One-Way ANOVA followed by Tukey’s multiple comparison test for significance. * stands for p < 0.05.

### TAT-pQYP restores response to capsaicin in PSNL mice by modulating expression coding for TRP channels

Since inhibition of TRPV1 had no effect on mechanical hypersensitivity assessed at 14 dpi, we tested the spontaneous nocifensive responses to capsaicin (CAP), a direct agonist of this channel, to assess if TRPV1 chemosensitivity activity was preserved. PSNL led to a decrease in nocifensive behavior latency compared to sham controls (not affected by TAT-pQYP treatment), which was modulated by TAT-pQYP (Fig. 3A, B), indicating a possible co-dependence of the lesion and PLC□ inhibition to elicit this phenotype. Since the PSNL group treated with vehicle control exhibited diminished response to CAP, we cannot affirm that the inhibitors are not modulating the response in this scenario, because the availability of TRPV1 could be compromised due to the injury. Thus, TAT-pQYP was administered 2.5 hours prior to CPZ injection to further confirm our hypothesis. In this case, CPZ significantly reduced the nocifensive behavior latency (Fig. 3C), suggesting that there could be an upregulation of TRPV1, induced by TAT-pQYP.

**Fig. 3:**
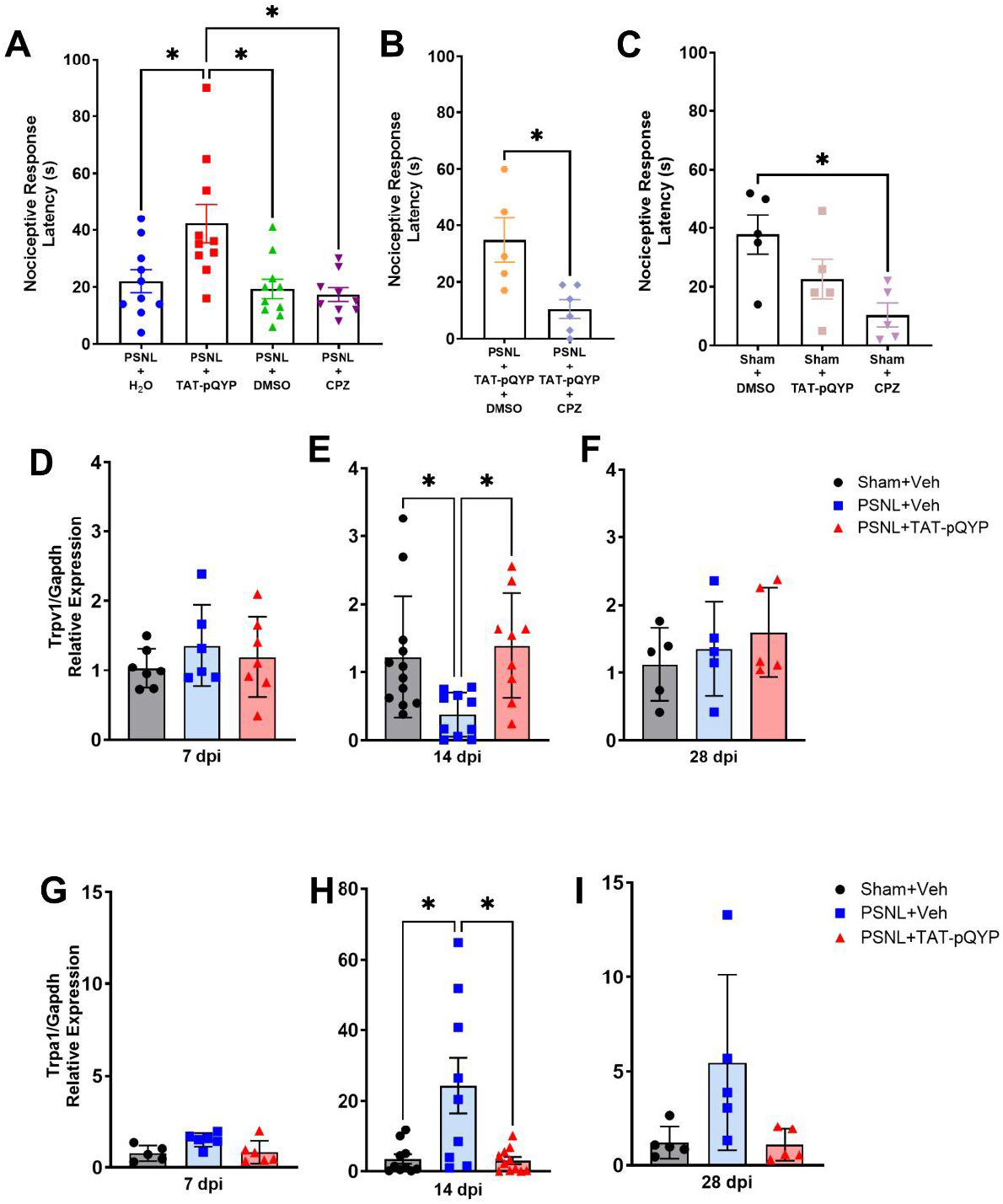
TAT-pQYP enhances capsaicin-evoked nocifensive behavior via upregulation of TRPV1. (A) TAT-pQYP enhances nocifensive behavior after capsaicin subcutaneous challenge, while capsazepine (CPZ) fails to modulate the response. (B) CPZ, but not TAT-pQYP, modulates sham operated mice nocifensive response to capsaicin. (C) CPZ treatment after TAT-pQYP reduces nocifensive response to capsaicin. (D-F) TRPV1 expression levels are not altered at 7 dpi (D) and 28 dpi (F), but are downregulated by PSNL and later normalized by TAT-pQYP treatment at 14 dpi (E). (G-I) TRPA1 expression levels are not altered at 7 dpi (G) and 28 dpi (I), but are upregulated by PSNL and later normalized by TAT-pQYP treatment at 14 dpi (H). Data are expressed as mean ± SD. One-Way ANOVA followed by Tukey’s multiple comparison test for significance (A, C-I), and unpaired t-test analysis for significance (B). * stands for p < 0.05.

Thus, to verify if alterations in channel activity corresponded to changes to their synthesis, we assessed the levels of TRPA1/V1 in the DRG. We observed that *Trpv1* was downregulated at the same time point, which was normalized to sham levels by TAT-pQYP (Figure 4D-F). Notably, *Trpa1* was significantly increased in the PSNL group only at 14 dpi, which was also normalized by TAT-pQYP (Fig. 3G-I). Altogether, these findings could explain both the lack of activity of CPZ regarding innocuous touch hypersensitivity (Fig. 2A) and the increased nocifensive behavior in response to CAP after TAT-pQYP treatment at 14 dpi (Fig. 3A).

**Fig. 4:**
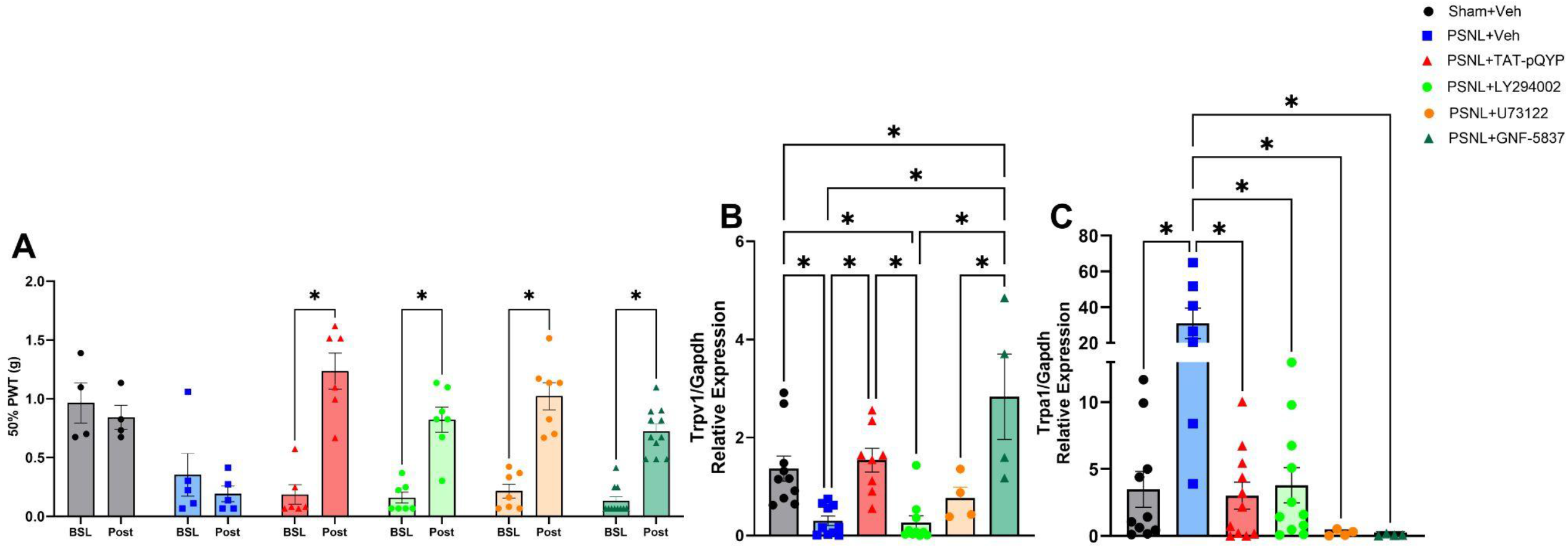
Modulation of Trk-PLC□-PI3K axis impacts mechanical hypersensitivity and normalizes TRP channels expression levels at 14 days post-injury. (A) TAT-pQYP, LY294002, U-73122 and GNF-5837 reverses mechanical hypersensitivity elicited by PSNL. (B) TAT-pQYP and GNF-5837, but not LY294002 or U-73122, normalizes TRPV1 expression to sham levels. (C) TAT-pQYP, LY294002, U-73122 and GNF-5837 reverses TRPA1 upregulation after PSNL. Data are expressed as mean ± SD. Two-Way ANOVA followed by Šídák’s multiple comparison test for significance (A), and one-way ANOVA followed by Tukey’s multiple comparison test for significance (B, C). * stands for p < 0.05.

### Mechanical hypersensitivity is dependent on the Trk-PLC axis, which influences TRP expression post-PSNL

To further substantiate the involvement of PLCγ signaling in the observed antinociceptive effect, we compared its effect over mechanical hypersensitivity with pan-Trk (GNF-5837) and pan-PLC (U-73122) inhibitors, and found comparable mechanical thresholds in these groups (Fig. 4A).

Of note, autophosphorylated Trk initiates other signaling cascades that have been described to modulate nociception, such as PI3K^40,41^. Thus, considering that Trk-PLC□ signaling modulates TRPV1 gating via hydrolysis of its inhibitory lipid PIP2^19^, while PI3K promotes PIP2 phosphorylation, we tested the effect of PI3K inhibition (LY294002) on the mechanical threshold. We found that LY294002, as TAT-pQYP, both reversed mechanical hypersensitivity (Fig. 4A). Furthermore, PI3K signaling has been described to be involved in transporting TRPV1 to the membrane^42^.

We also compared Trk-PLC-PI3K signaling axis modulators in terms of TRP channels expression. We found that *Trpv1* was upregulated by PLC□ (TAT-pQYP) and Trk (GNF-5837) inhibition, while PI3K (LY294002) inhibition did not alter its expression profile (Fig. 4B). However, all modulators were able to downregulate *Trpa1* expression (Fig. 4C), suggesting differential regulatory mechanisms of expression of TRP channels via Trk signaling. Of note, the pan-PLC inhibitor failed to mimic the TAT-pQYP effect over TRPV1, even though it downregulated TRPA1.

## DISCUSSION

Neuropathic pain remains a major clinical challenge due to its complex pathophysiology and the limited efficacy of currently available treatments. TRPA1/V1 channels have been widely explored as potential targets to alleviate pain, since they have enhanced expression and activity during inflammatory and neuropathic pain, and are often upregulated in peripheral somatosensory neurons^43–45^. Many knockout studies have reported mitigated mechanical and thermal hypersensitivity in a variety of pain models, supporting the fact that direct and specific inhibition of these channels can be a good alternative for pain management^40,46–48^. However, clinical trials have been challenging, and suggest that such strategy can lead to multiple side effects or lack efficacy^49^, for instance, capsaicin-based patches promoting TRPV1 dessensitization have been associated with reduced local pain thresholds, but lack efficiency in diffuse pain and affects heat sensitivity^50,51^.

Previous studies have suggested that interfering with TRP channel activity by targeting signaling cascades and key protein-protein interaction with the channels leading to channel activation (as reviewed in: ^52^), such as interfering with TRPV1-AKAP79 (A-Kinase Anchoring Protein 79)^53^, or mimicking TRPV1-KChIP3 (Potassium voltage-gated channel interacting protein 3), both which successfully mitigated inflammatory pain in mice^54^. Here, we provide compelling evidence that PLCγ is a critical mediator of peripheral sensitization in neuropathic pain, acting by modulating TRPV1 and TRPA1 during PSNL. By employing TAT-pQYP, designed to disrupt the interaction between PLCγ and RTKs, we observed a significant reversal of mechanical hypersensitivity and molecular alterations consistent with antinociceptive effects.

Regarding mRNA expression, we observed a dynamic regulation of the channels related to both the time post-injury and TAT-pQYP treatment, inhibiting PLCγ mediated signaling. TRPA1 presented a characteristic upregulation profile, peaking at day 14 after surgery, matching the mechanical hypersensitivity profile observed. In contrast, TRPV1 levels were downregulated at 14 dpi and similar to sham levels, at 7 and 28 dpi. TAT-pQYP treatment affected both expression level and nociception. A study involving trigeminal nerve injury and resiniferatoxin-induced pain^24,25^ reported downregulation of TRPV1, mediated by neuron loss after injury. We can not discard that at 14 dpi a possible TRPV1^+^ neuron loss is impacting our results, however we were able to restore TRPV1 expression upon TAT-pQYP treatment, indicating that there are still viable TRPV1^+^ neurons.

A previous report showed that while capsazepine was able to reverse carrageenan- and capsaicin-induced hypersensitivity, it failed to impact PSNL-induced hypersensitivity in mice and rats, despite showing efficacy in guinea pigs^55^, pointing to species differences in pharmacological targeting of the channel. Here, we observed that capsazepine was effective to reverse mechanical hypersensitivity at 28, but not 14 dpi, adding that TRPV1 direct targeting depends on the progression of the neuropathy to be pharmacologically relevant. On the other hand, TRPA1 inhibition by HC-030031 successfully reversed mechanical hypersensitivity in both timepoints, as reported by others^56,57^.

We recently described that TAT-pQYP was not able to interfere with TRPV1 chemosensitivity to capsaicin in naive conditions, but was effective in disrupting TrkA-PLC□-TRPV1 signaling triggered by inflammation, promoting antinociception and decrease in edema via PKC activation^13^. Here, TAT-pQYP did not interfere with TRPV1 activity or expression in sham operated animals and did not increase channel levels at 7 or 28 dpi, probably due to lack of growth factor mediated signaling, while hypersensitivity of PSNL animals was reversed at all timepoints and TRP channel expression at 14 dpi was normalized to sham levels.

Understanding the molecular mechanisms that regulate TRP channel activity is key for the development of improved analgesics. We show evidence that targeting TRPV1 may not be a viable option for pain management during the entire time course of neuropathy, and that during PSNL, the hypersensitive state is established by TRPA1 when TRPV1 is downregulated. TRPV1/A1 often act together to regulate nociception^26,58^ and have been explored in the context of the heteromer in which TRPV1 tightly regulates TRPA1 gating. The transmembrane protein 100 (Tmem100), which weakens the association between TRPV1 and TRPA1 by binding to both. In fact, mice lacking Tmem100 have reduced TRPA1-mediated pain, without interfering with TRPV1-mediated pain^59^.

Considering that when co-expressed in the heteromer form, TRPA1 activity is inhibited by TRPV1 in the closed state^59,60^, and that TAT-pQYP prevents PIP2 hydrolysis by PLC□, sustaining TRPV1 inhibition, could explain how the peptide acts on both channels when they are both expressed (7 and 28 dpi). Furthermore, the downregulation of TRPA1 expression by TAT-pQYP may also be an important mechanism for reversal of nociception since TRPV1 expression is damped at 14 dpi, and TRPA1 levels are exceedingly upregulated sustaining hypersensitivity, besides PIP2 inhibiting the channel^20^.

The exact mechanism by which disrupting RTK-PLC□ interaction during neuropathic pain downregulates TRPA1 and enhances TRPV1 should be further investigated, since expression level increased at three hours post treatment with TAT-pQYP, implying that PLC□ could be impacting on the expression of TRPA1 and that the decrease in TRPA1 could lead to an increase in TRPV1.

The limitations of this study are the sole use of male mice and that TAT-pQYP could have other targets besides PLCγ. Taken together we show that PLCγ is a key player in PSNL induced mechanical hypersensitivity and that it could be targeted to mitigate neuropathic pain.

## Acknowledgments

Dr. Élora Midavaine for critically reading the manuscript. Dr. Vanessa O. Zambelli for donating reagents. Luciana Silva for general technical assistance.

## Data availability

The authors declare that all the data supporting the findings of this study are contained within the paper.

## Author contributions

D.S. and C.S.D. acquired funding and supervised the research; D.S., C.S.D., B.C.M. and A.M.d.N. conceived and designed experiments; B.C.M., A.M.d.N., E.F.T., A.M.R., C.P.G. performed experiments; B.C.M., A.M.d.N. and D.S. analyzed/interpreted the data and wrote the manuscript; All authors revised and approved the final version of the manuscript.

## Funding

This work was supported by FAPESP (Fundação de Amparo à Pesquisa do Estado de São Paulo): Grant No. 2019/06982-6 (to D.S. and C.S.D.), fellowship No. 2022/00594-7 (to B.C.M.), fellowship No. 24/18174-0 (to A.M.d.N.), fellowship No. 2023/10572-3 (to A.M.R.), fellowship No. 2020/16204-8 (C.P.G); CNPq (Conselho Nacional de Desenvolvimento Científico e Tecnológico): fellowship PQ-2 No. 307739/2018-0 (to D.S.).

## SUPPLEMENTARY DATA FILE LEGENDS

**DATA S1:** Details on all statistical tests applied for analysis. Data include type of test, post hoc test, sample space (N), degrees of freedom, statistics summary, mean differences, 95% confidence intervals and exact p-values.

## Notes

### Competing Interest Statement

The authors have declared no competing interest.

